## Supplementary Materials for "Human brain representations of internally generated outcomes of approximate calculation revealed by ultra-high-field brain imaging"

Supplementary Methods:

To explore effects of the different experimental effects of interest (sample numerosity, operation, operand, result numerosity) on brain activity in univariate analyses using standard models of the hemodynamic response, an additional first level model of the fMRI data was computed in each subject. This model included the following set of predictors for the standard trial types (catch trials being modelled separately and not included in the analysis): 4 predictors for the 4 different sample numerosities (6, 12, 24, 48) with stimulus onset at the time of sample display appearance, 2 predictors for operands (2 and 4), timed at the onset of the operation instruction display, two predictors for the operations (multiplication and division) with their onset defined at 500 ms after the operation instruction display, 4 predictors for the possible result numerosities (6, 12, 24, 48) with their onset defined at 1s after the operation instruction display, and 8 predictors for the match numerosity (separated into smaller and larger for each of the four results) at the onset of the match display. All predictor were modelled as stick functions convolved with the canonical hemodynamic response function of SPM12. The GLM included the six motion parameters as covariate of no interest, an AR(1) model was used to account for serial auto-correlation and low-frequency signal drifts were removed by a high-pass filter with a cutoff of 244 s. The following contrasts were created at the individual subject level: The parametric increase with numerosity, separately for the sample and result numerosity (contrast weights -1.5 -0.5 0.5 1.5) and differential contrasts for each individual numerosity vs all others, separately for the sample and result (e.g., contrast weights [3 -1 -1 -1] for numerosity 6). For each of these, second level analyses were then conducted using one-sample t-tests in Freesurfer (after projecting all contrasts into FsAverage space). Results are presented in Supplementary Figure 1 and Supplementary Table 1.

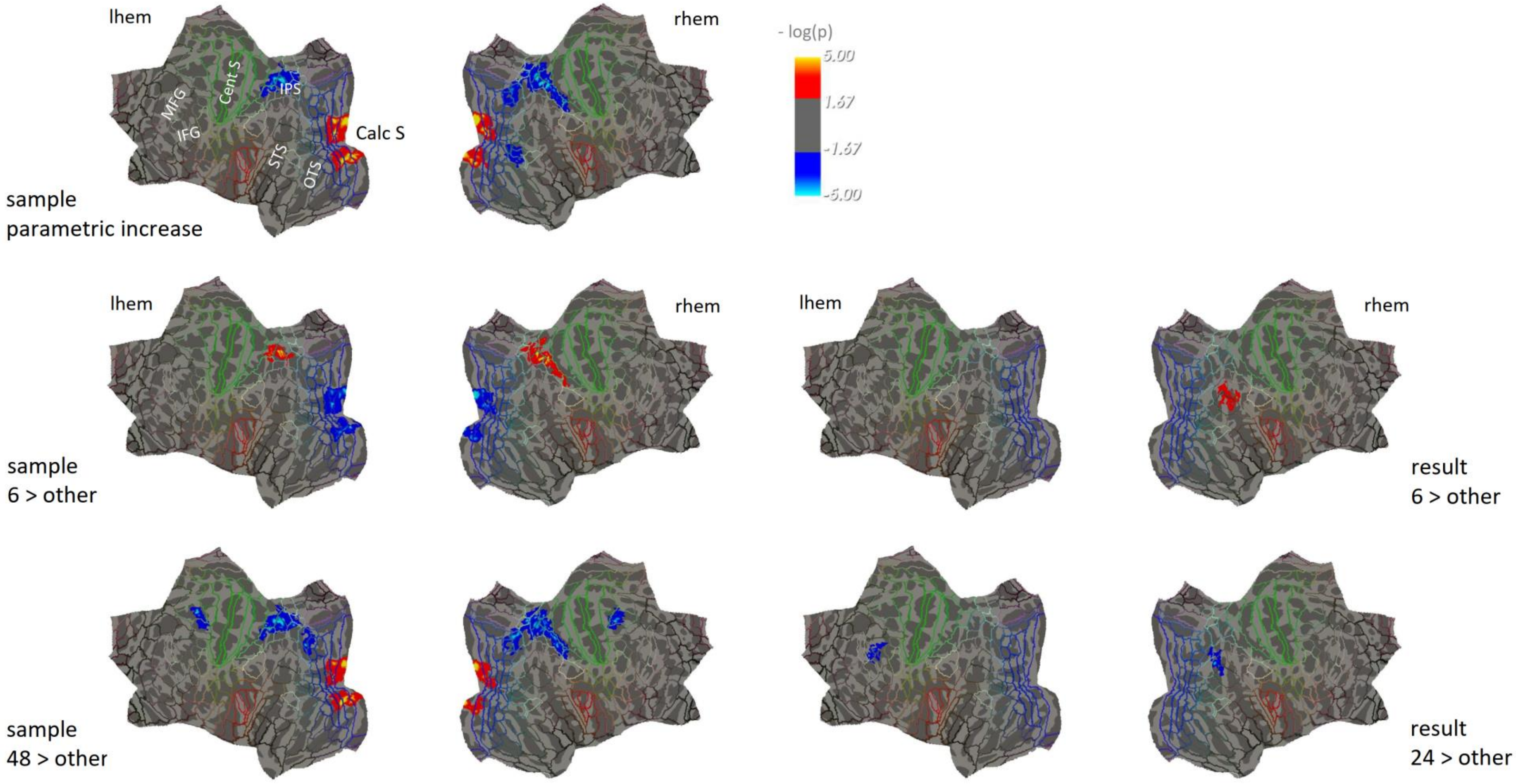


*Supplementary Figure 1 : Results of a random effects group analysis (n = 17) testing for parametric increases with numerosity or differential effects of each specific numerosity vs all others on univariate brain responses. Surface-based group analyses were performed in FreeSurfer (one-sample t-tests, correction for multiple comparisons by permutation at cluster level, pFWE <.05, cluster forming threshold p < .01). The color scale represents uncorrected significance within the clusters surviving correction. Results are projected onto a flattened cortical average surface (https://mri.sdsu.edu/sereno/csurf/), with superimposed lines representing the regional borders of the HCP-MMP1 parcellation.*

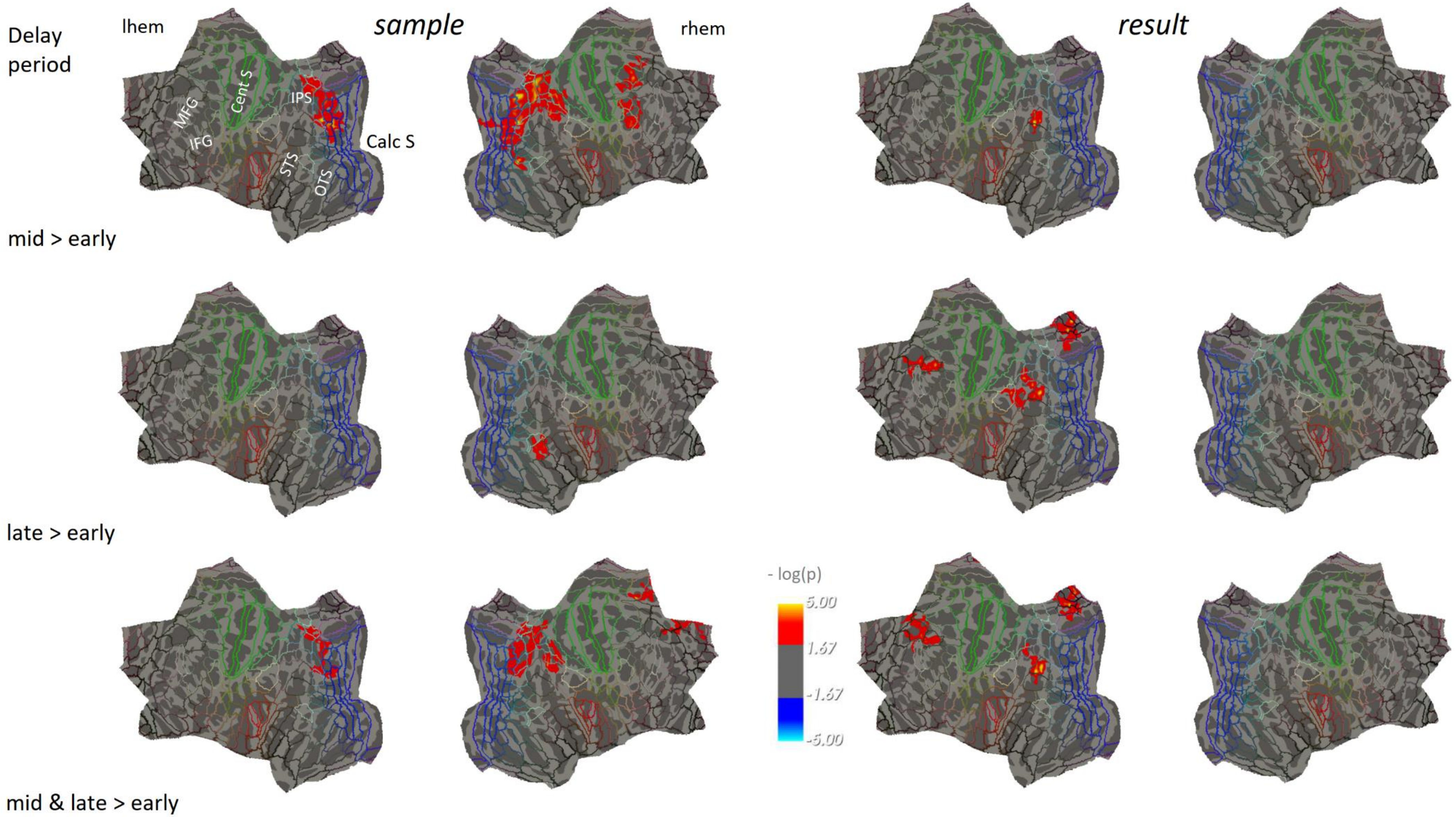


*Supplementary Figure 2 : Results of a searchlight analysis (n = 17) testing for differential effects between time windows in multiple regression-based representational similarity analysis. Surface-based group analyses were conducted in FreeSurfer (one-sample t-tests, correction for multiple comparisons by permutation at cluster level, pFWE <.05, cluster forming threshold p < .01). The color scale represents uncorrected significance within the clusters surviving correction. Results are projected onto a flattened cortical average surface (https://mri.sdsu.edu/sereno/csurf/), with superimposed lines representing the regional borders of the HCP-MMP1 parcellation.*

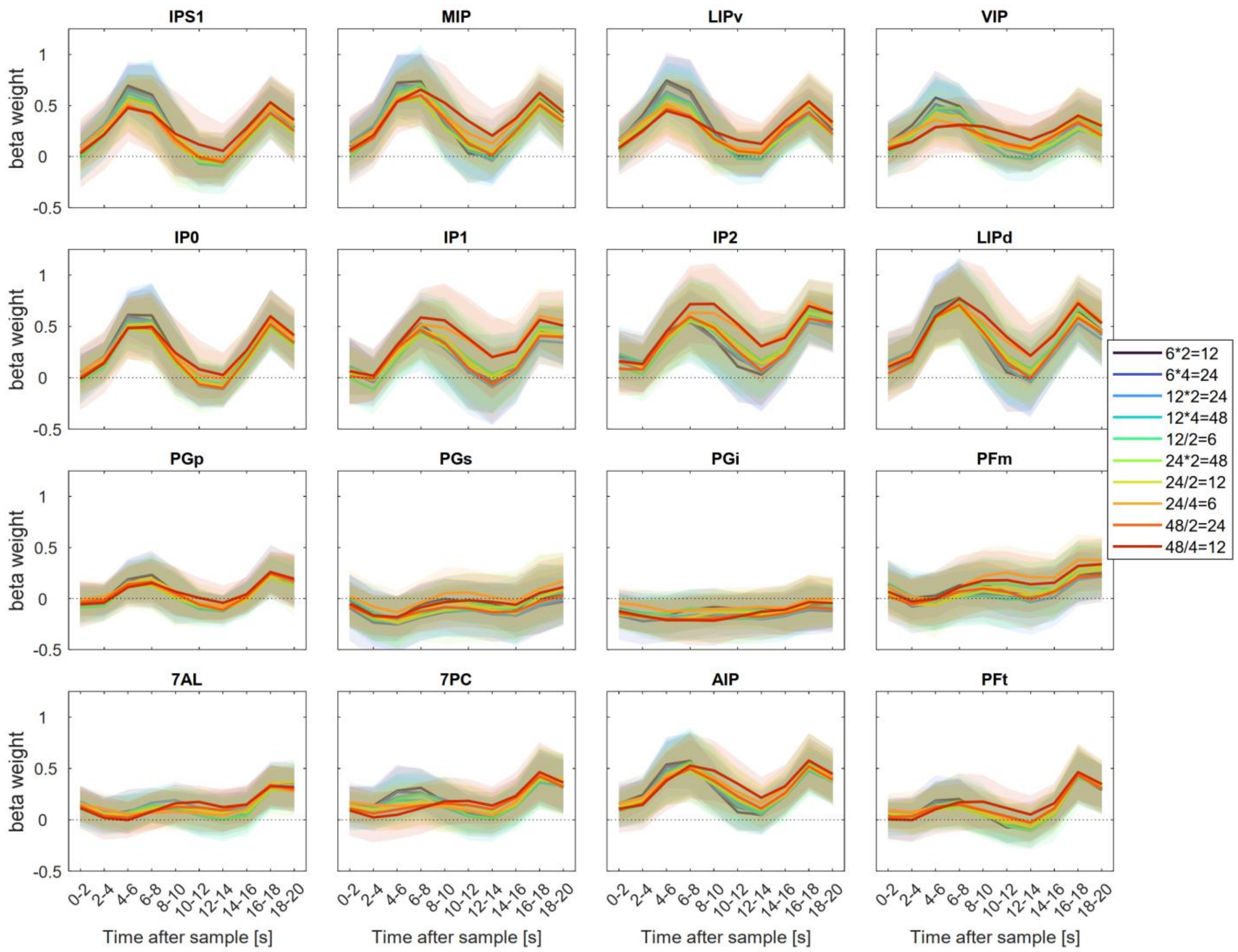


*Supplementary Figure 3 : Evoked activity time courses as computed by FIR estimates for all parietal regions of interest for the 10 experimental conditions (N=17, means and standard deviations across subjects).*

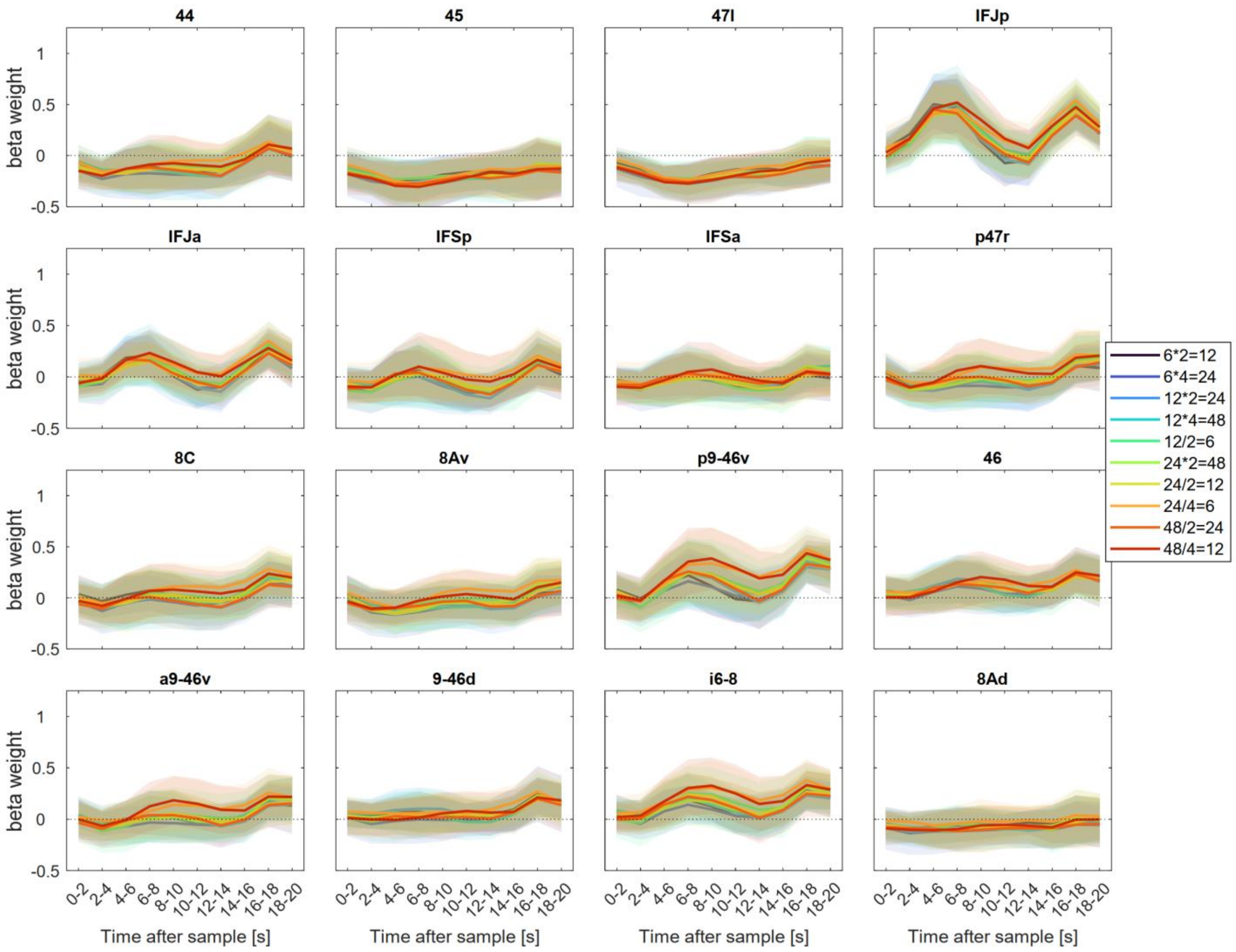


*Supplementary Figure 4 : Evoked activity time courses as computed by FIR estimates for all parietal regions of interest for the 10 experimental conditions (N=17, means and standard deviations across subjects).*

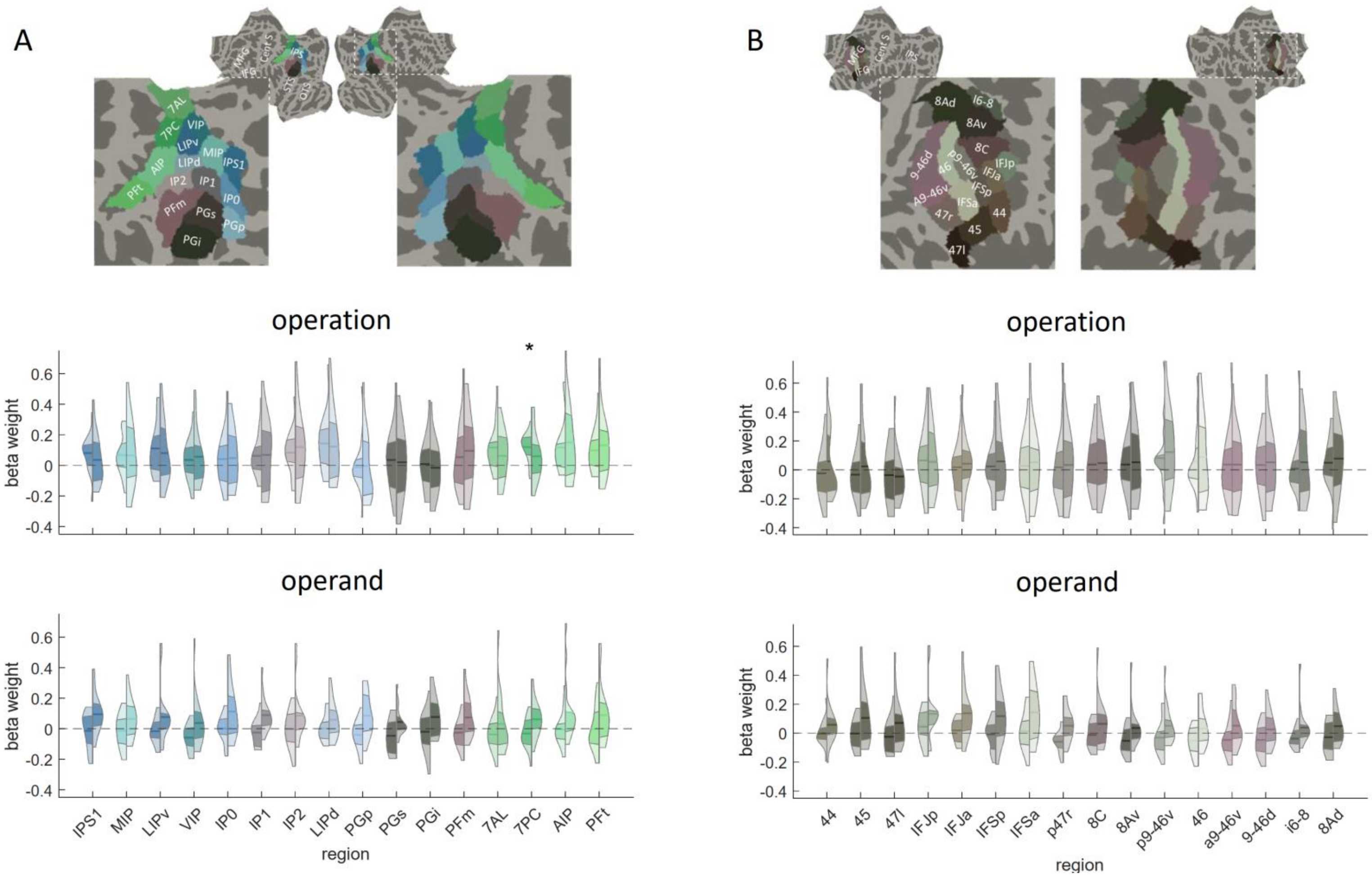


*Supplementary Figure 5 : Results of multiple regression-based representational similarity analysis for the effects of operation and operand predictors on brain activity patterns within subregions of parietal (A) and lateral prefrontal (B) cortex. ROI delineation is based on the HCP_MMP1 parcellation shown here on the flattened cortical average surface. For each ROI, violin plots show the distribution of beta estimates for operation and operand predictors (n = 17). The central horizontal line indicates the mean, and the adjacent colored area the limits of the first and third quartile, with the left part of the violin for each ROI corresponding to averages of the middle (4-8 s), and the right part to averages of the late (8-12 s) delay period. Stars indicate significantly larger responses than 0 in two-tailed t-tests, after FDR correction for multiple comparisons (pFDR<.05, across all 32 subregions and the two time periods).*

**Supplementary Tables**

**Supplementary Table 1:** Cluster summary table for a surface-based group analysis (N=17) conducted on univariate effects of sample and result numerosity when modelling responses by a canonical HRF.

For each contrast displayed in Supplementary Figure 2, and each cluster surviving $p_{FWE} < .05$ (corrected at cluster level by permutation methods with cluster forming threshold p < .01) the table reports: the cluster label for the activation maximum (as defined by the anatomical labels from the Destrieux Atlas), the maximum -log10(p) value in the cluster (Max), the cluster surface area in mm2 (Size), the MNI coordinates of the maximally activated vertex within each cluster (MNI X, Y, Z), the cluster-wise p-value of each cluster (CWP), the number of vertices included in each cluster (NVtxs).

| **Sample parametric increase** | | | | | | | |
|---|---|---|---|---|---|---|---|
| *Left hemisphere* | | | | | | | |
| Cluster Label | Max | Size (mm2) | MNI X | MNI Y | MNI Z | CWP | NVtxs |
| G_oc-temp_med-Lingual | 7.08 | 2416.5 | -8.1 | -93.3 | -10.2 | 0.0004 | 2883 |
| G_parietal_sup | -6.86 | 1395.1 | -29.4 | -55.7 | 56 | 0.0060 | 3253 |
| *Right hemisphere* | | | | | | | |
| Cluster Label | Max | Size (mm2) | MNI X | MNI Y | MNI Z | CWP | NVtxs |
| S_intrapariet_and_P_trans | -6.21 | 2804.6 | 34.5 | -46.4 | 49.1 | 0.0008 | 6100 |
| Pole_occipital | 7.62 | 1936.9 | 14.4 | -97.4 | 4.3 | 0.0024 | 2341 |
| G_and_S_occipital_inf | -3.59 | 637.4 | 45.4 | -64.3 | -11.3 | 0.0290 | 874 |
| **Sample 6 > other** | | | | | | | |
| *Left hemisphere* | | | | | | | |
| Cluster Label | Max | Size (mm2) | MNI X | MNI Y | MNI Z | CWP | NVtxs |
| G_oc-temp_med-Lingual | -6.36 | 2078.2 | -8.1 | -92.8 | -10.4 | 0.0024 | 2461 |
| G_parietal_sup | 5.71 | 696.6 | -28.9 | -55.7 | 55.4 | 0.0183 | 1598 |
| *Right hemisphere* | | | | | | | |
| Cluster Label | Max | Size (mm2) | MNI X | MNI Y | MNI Z | CWP | NVtxs |
| Pole_occipital | -5.97 | 1599.8 | 22.2 | -99.5 | 0.3 | 0.0028 | 1952 |
| S_intrapariet_and_P_trans | 6.21 | 1511.9 | 25.4 | -55.5 | 56.7 | 0.0036 | 3568 |
| **Sample 48 > other** | | | | | | | |
| *Left hemisphere* | | | | | | | |
| Cluster Label | Max | Size (mm2) | MNI X | MNI Y | MNI Z | CWP | NVtxs |
| G_oc-temp_med-Lingual | 6.29 | 2135.2 | -17.9 | -86.1 | -9.5 | 0.0016 | 2516 |
| G_parietal_sup | -6.68 | 1481.1 | -30.1 | -55.6 | 56.3 | 0.0024 | 3458 |
| S_precentral-sup-part | -5.97 | 420.5 | -22.5 | -8.4 | 48.2 | 0.0400 | 893 |
| S_intrapariet_and_P_trans | -5.34 | 407.9 | -25.4 | -70.3 | 23 | 0.0423 | 747 |
| *Right hemisphere* | | | | | | | |

| Cluster Label | Max | Size (mm2) | MNI X | MNI Y | MNI Z | CWP | NVtxs |
|---|---|---|---|---|---|---|---|
| S_intrapariet_and_P_trans | -6.17 | 2666.2 | 25.3 | -58.8 | 47.7 | 0.0004 | 5834 |
| Pole_occipital | 5.59 | 1404.7 | 14.8 | -94.8 | 4.7 | 0.0044 | 1713 |
| S_precentral-sup-part | -5.17 | 458.3 | 31.8 | -9.7 | 48.9 | 0.0373 | 1056 |
| **Result 6 > other** | | | | | | | |
| *Right hemisphere* | | | | | | | |
| Cluster Label | Max | Size (mm2) | MNI X | MNI Y | MNI Z | CWP | NVtxs |
| G_pariet_inf-Angular | 5.15 | 766.5 | 53.2 | -52.1 | 30.8 | 0.01196 | 1568 |
| **Result 24 > other** | | | | | | | |
| *Left hemisphere* | | | | | | | |
| Cluster Label | Max | Size (mm2) | MNI X | MNI Y | MNI Z | CWP | NVtxs |
| S_front_inf | -4.60 | 543.3 | -40.2 | 21.5 | 30.2 | 0.0262 | 1006 |
| *Right hemisphere* | | | | | | | |
| Cluster Label | Max | Size (mm2) | MNI X | MNI Y | MNI Z | CWP | NVtxs |
| G_pariet_inf-Angular | -5.08 | 541.8 | 38.7 | -73.6 | 33.3 | 0.0317 | 902 |

**Supplementary Table 2:** Cluster summary table for differential contrasts between different parts of the delay period for the surface-based group analysis (N=17) conducted on the searchlight pattern analysis (multiple regression with sample, operation, operand and result predictors on the fMRI distance matrices).

For each contrast displayed in Supplementary Figure 2, and each cluster surviving $p_{FWE} < .05$ (corrected at cluster level by permutation methods with cluster forming threshold $p < .01$) the table reports: the cluster label for the activation maximum (as defined by the anatomical labels from the Destrieux Atlas), the maximum -log10(p) value in the cluster (Max), the cluster surface area in mm2 (Size), the MNI coordinates of the maximally activated vertex within each cluster (MNI X, Y, Z), the cluster-wise p-value of each cluster (CWP), the number of vertices included in each cluster (NVtxs). No above threshold clusters were detected for the operation and the operand at any time window.

| **Sample – middle > early delay** | | | | | | | |
|---|---|---|---|---|---|---|---|
| *Left hemisphere* | | | | | | | |
| Cluster Label | Max | Size (mm2) | MNI X | MNI Y | MNI Z | CWP | NVtxs |
| S_oc_sup_and_transversal | 5.55 | 2700.1 | -34.3 | -79.8 | 18.4 | 0.0020 | 4573 |
| *Right hemisphere* | | | | | | | |
| Cluster Label | Max | Size (mm2) | MNI X | MNI Y | MNI Z | CWP | NVtxs |
| G_occipital_middle | 5.96 | 5932.8 | 36.5 | -82.5 | 20.9 | 0.0004 | 11367 |
| S_front_sup | 4.57 | 910.7 | 27.4 | 1.7 | 48.9 | 0.0243 | 1745 |
| S_precentral-inf-part | 4.02 | 878.1 | 49.0 | 9.9 | 12.2 | 0.0270 | 1674 |
| **Sample – late > early delay** | | | | | | | |

| Right hemisphere | | | | | | | |
|---|---|---|---|---|---|---|---|
| Cluster Label | Max | Size (mm2) | MNI X | MNI Y | MNI Z | CWP | NVtxs |
| G_temporal_middle | 4.78 | 669.8 | 61.1 | -46.9 | 0.5 | 0.0459 | 1157 |
| **Sample – late > middle delay** | | | | | | | |
| *Right hemisphere* | | | | | | | |
| Cluster Label | Max | Size (mm2) | MNI X | MNI Y | MNI Z | CWP | NVtxs |
| G_parietal_sup | -4.24 | 503.9 | 26.6 | -54.8 | 57.7 | 0.0490 | 1109 |
| **Sample – late and middle > early delay** | | | | | | | |
| *Left hemisphere* | | | | | | | |
| Cluster Label | Max | Size (mm2) | MNI X | MNI Y | MNI Z | CWP | NVtxs |
| G_occipital_sup | 4.00 | 1249.6 | -21.5 | -80.5 | 33.1 | 0.0183 | 2194 |
| *Right hemisphere* | | | | | | | |
| Cluster Label | Max | Size (mm2) | MNI X | MNI Y | MNI Z | CWP | NVtxs |
| S_intrapariet_and_P_trans | 4.15 | 2082.8 | 20.2 | -59.9 | 52.0 | 0.0060 | 3956 |
| G_pariet_inf-Angular | 4.49 | 852.7 | 46.7 | -53.7 | 44.0 | 0.0317 | 2123 |
| G_front_sup | 5.02 | 846.7 | 9.1 | 11.7 | 56.4 | 0.0321 | 1689 |
| **Result – middle > early delay** | | | | | | | |
| *Left hemisphere* | | | | | | | |
| Cluster Label | Max | Size (mm2) | MNI X | MNI Y | MNI Z | CWP | NVtxs |
| S_temporal_sup | 5.14 | 459.2 | -40.1 | -63.9 | 33.0 | 0.0388 | 919 |
| **Result – late > early delay** | | | | | | | |
| *Left hemisphere* | | | | | | | |
| Cluster Label | Max | Size (mm2) | MNI X | MNI Y | MNI Z | CWP | NVtxs |
| G_pariet_inf-Angular | 5.11 | 1543.5 | -37.9 | -72.5 | 36.4 | 0.0120 | 3423 |
| S_subparietal | 5.02 | 872.9 | -7.4 | -48.8 | 29.6 | 0.0373 | 1812 |
| G_front_middle | 4.56 | 794.5 | -40.1 | 10.9 | 50.7 | 0.0463 | 1398 |
| **Result – middle and late > early delay** | | | | | | | |
| *Left hemisphere* | | | | | | | |
| Cluster Label | Max | Size (mm2) | MNI X | MNI Y | MNI Z | CWP | NVtxs |
| S_front_sup | 4.20 | 1020.0 | -27.1 | 31.9 | 28.1 | 0.0207 | 1760 |
| S_subparietal | 5.04 | 881.7 | -10.5 | -57.2 | 29.9 | 0.0258 | 1828 |
| S_temporal_sup | 5.40 | 822.3 | -40.3 | -63.6 | 33.4 | 0.0298 | 1729 |

**Supplementary Table 3:** Results (N=17) of cross-decoding performance between sample and result numerosity representations for the 16 parietal and 16 frontal ROIs derived from the HCP-MMP1 parcellation. For each ROI, the mean of the decoding performance corresponding to a correlation score (mean) is reported together with effect size (Cohen's d), t-value of two-tailed t-tests against 0, degrees of freedom, uncorrected p values, lower and upper bound of the 95% confidence interval around the mean, and corrected p values. Significance was corrected for multiple comparisons across all ROIs by false discovery rate (FDR).

| ROI | mean | Cohen's d | t | df | $p_{uncorr}$ | $CI_l$ | $CI_u$ | $p_{FDR\ corr}$ |
|---|---|---|---|---|---|---|---|---|
| IPS1 | 0.043 | 0.433 | 1.88 | 16 | 0.0789 | 0.003 | 0.084 | 0.6315 |
| MIP | 0.055 | 0.593 | 2.57 | 16 | 0.0206 | 0.018 | 0.092 | 0.5580 |
| LIPv | 0.030 | 0.296 | 1.28 | 16 | 0.2188 | -0.011 | 0.071 | 0.7862 |
| VIP | 0.017 | 0.214 | 0.93 | 16 | 0.3685 | -0.015 | 0.050 | 0.7862 |
| IP0 | 0.038 | 0.484 | 2.10 | 16 | 0.0523 | 0.006 | 0.069 | 0.5580 |
| IP1 | 0.017 | 0.228 | 0.99 | 16 | 0.3389 | -0.013 | 0.047 | 0.7862 |
| IP2 | 0.004 | 0.046 | 0.20 | 16 | 0.8454 | -0.033 | 0.042 | 0.9360 |
| LIPd | 0.006 | 0.069 | 0.30 | 16 | 0.7674 | -0.028 | 0.039 | 0.9360 |
| PGp | 0.019 | 0.335 | 1.45 | 16 | 0.1664 | -0.004 | 0.043 | 0.7605 |
| PGs | 0.012 | 0.117 | 0.51 | 16 | 0.6195 | -0.030 | 0.055 | 0.9360 |
| PGi | 0.024 | 0.218 | 0.94 | 16 | 0.3596 | -0.020 | 0.068 | 0.7862 |
| PFm | -0.006 | -0.063 | 0.27 | 16 | 0.7875 | -0.041 | 0.030 | 0.9360 |
| 7AL | 0.015 | 0.166 | 0.72 | 16 | 0.4837 | -0.022 | 0.052 | 0.9104 |
| 7PC | 0.005 | 0.061 | 0.27 | 16 | 0.7940 | -0.028 | 0.038 | 0.9360 |
| AIP | 0.021 | 0.235 | 1.02 | 16 | 0.3243 | -0.015 | 0.057 | 0.7862 |
| PFt | 0.020 | 0.231 | 1.00 | 16 | 0.3315 | -0.015 | 0.056 | 0.7862 |
| 44 | 0.018 | 0.231 | 1.00 | 16 | 0.3323 | -0.014 | 0.050 | 0.7862 |
| 45 | 0.003 | 0.035 | 0.15 | 16 | 0.8817 | -0.027 | 0.032 | 0.9360 |
| 47l | 0.012 | 0.154 | 0.67 | 16 | 0.5145 | -0.019 | 0.042 | 0.9146 |
| IFJp | 0.022 | 0.377 | 1.63 | 16 | 0.1219 | -0.002 | 0.046 | 0.6500 |
| IFJa | -0.005 | -0.095 | 0.41 | 16 | 0.6867 | -0.027 | 0.017 | 0.9360 |
| IFSp | 0.026 | 0.381 | 1.65 | 16 | 0.1189 | -0.002 | 0.053 | 0.6500 |
| IFSa | 0.008 | 0.141 | 0.61 | 16 | 0.5512 | -0.015 | 0.031 | 0.9283 |
| p47r | 0.018 | 0.266 | 1.15 | 16 | 0.2655 | -0.009 | 0.046 | 0.7862 |
| 8C | 0.001 | 0.007 | 0.03 | 16 | 0.9772 | -0.036 | 0.037 | 0.9772 |
| 8Av | 0.002 | 0.028 | 0.12 | 16 | 0.9038 | -0.027 | 0.031 | 0.9360 |
| p9-46v | 0.004 | 0.050 | 0.22 | 16 | 0.8303 | -0.027 | 0.035 | 0.9360 |
| 46 | 0.010 | 0.126 | 0.55 | 16 | 0.5923 | -0.023 | 0.044 | 0.9360 |
| a9-46v | -0.002 | -0.027 | 0.12 | 16 | 0.9068 | -0.037 | 0.032 | 0.9360 |
| 9-46d | -0.012 | -0.198 | 0.86 | 16 | 0.4042 | -0.035 | 0.012 | 0.8084 |
| i6-8 | 0.036 | 0.499 | 2.16 | 16 | 0.0463 | 0.007 | 0.065 | 0.5580 |
| 8Ad | -0.004 | -0.039 | 0.17 | 16 | 0.8682 | -0.041 | 0.034 | 0.9360 |

**Supplementary Table 4:** Results (N=17) of the correlation analysis between cross-decoding performance between sample and result numerosity representations and behavioral Weber fractions, for the 16 parietal and 16 frontal ROIs derived from the HCP-MMP1 parcellation. For each ROI, the Pearson correlation coefficient (r) is reported together with uncorrected as well as corrected p values, and the lower and upper bound of the 95% confidence interval of the correlation coefficient. Significance was corrected for multiple comparisons across all ROIs by false discovery rate (FDR).

| **ROI** | **r** | **$p_{uncorr}$** | **$p_{FDR corr}$** | **$CI_l$** | **$CI_u$** | **ROI** | **rho** | **$p_{uncorr}$** | **$CI_l$** | **$CI_u$** | **$p_{FDR corr}$** |
|---|---|---|---|---|---|---|---|---|---|---|---|
| IPS1 | -0.23 | 0.3852 | 0.8632 | -0.64 | 0.29 | 44 | 0.29 | 0.2585 | -0.22 | 0.68 | 0.8632 |
| MIP | -0.80 | 0.0001 | 0.0035 | -0.93 | -0.52 | 45 | -0.16 | 0.5398 | -0.59 | 0.35 | 0.8632 |
| LIPv | -0.37 | 0.1442 | 0.8632 | -0.72 | 0.13 | 47l | -0.22 | 0.3875 | -0.64 | 0.29 | 0.8632 |
| VIP | -0.06 | 0.8190 | 0.9719 | -0.53 | 0.43 | IFJp | -0.16 | 0.5326 | -0.60 | 0.34 | 0.8632 |
| IP0 | -0.18 | 0.5006 | 0.8632 | -0.61 | 0.33 | IFJa | -0.17 | 0.5045 | -0.60 | 0.33 | 0.8632 |
| IP1 | -0.12 | 0.6474 | 0.8632 | -0.57 | 0.38 | IFSp | -0.23 | 0.3649 | -0.64 | 0.28 | 0.8632 |
| IP2 | -0.06 | 0.8272 | 0.9719 | -0.52 | 0.44 | IFSa | -0.12 | 0.6415 | -0.57 | 0.38 | 0.8632 |
| LIPd | -0.13 | 0.6063 | 0.8632 | -0.58 | 0.37 | p47r | -0.46 | 0.0647 | -0.77 | 0.03 | 0.8632 |
| PGp | -0.05 | 0.8576 | 0.9719 | -0.52 | 0.44 | 8C | 0.01 | 0.9583 | -0.47 | 0.49 | 0.9875 |
| PGs | -0.36 | 0.1515 | 0.8632 | -0.72 | 0.14 | 8Av | -0.22 | 0.3910 | -0.64 | 0.29 | 0.8632 |
| PGi | -0.23 | 0.3746 | 0.8632 | -0.64 | 0.28 | p9-46v | 0.25 | 0.3390 | -0.27 | 0.65 | 0.8632 |
| PFm | -0.25 | 0.3253 | 0.8632 | -0.65 | 0.26 | 46 | -0.08 | 0.7625 | -0.54 | 0.42 | 0.9719 |
| 7AL | -0.17 | 0.5140 | 0.8632 | -0.60 | 0.34 | a9-46v | -0.25 | 0.3362 | -0.65 | 0.26 | 0.8632 |
| 7PC | -0.19 | 0.4637 | 0.8632 | -0.61 | 0.32 | 9-46d | -0.12 | 0.6432 | -0.57 | 0.38 | 0.8632 |
| AIP | 0.04 | 0.8808 | 0.9719 | -0.45 | 0.51 | i6-8 | 0.00 | 0.9875 | -0.48 | 0.48 | 0.9875 |
| PFt | -0.19 | 0.4671 | 0.8632 | -0.61 | 0.32 | 8Ad | -0.02 | 0.9429 | -0.49 | 0.47 | 0.9875 |
